# Sex and Disease Regulate MHC I Expression in Human Lung Epithelial Cells

**DOI:** 10.1101/2024.06.20.599895

**Authors:** Justine Mathé, Sylvie Brochu, Damien Adam, Emmanuelle Brochiero, Claude Perreault

**Affiliations:** Institute for Research in Immunology and Cancer; Département de Médecine, Université de Montréal, Montréal, Québec, Canada; Centre de Recherche du CHUM (CRCHUM), Montréal, Québec, Canada

**Keywords:** Antigen presentation, MHC I, Lung epithelial cells, Chronic respiratory diseases, Sexual dimorphism

## Abstract

Major histocompatibility complex class I (MHC I) molecules present endogenous peptides to CD8+ T-cells for immunosurveillance of infections and cancers. Recent studies revealed unexpected heterogeneity in MHC I expression among cells of different lineages. While respiratory diseases rank among the leading causes of mortality, studies in mice showed that lung epithelial cells (LECs) express lower MHC I levels than all other tested cell types. The present study aimed to evaluate MHC I expression in human LECs from parenchymal explants using single-cell RNA sequencing (scRNA-seq) and immunostaining of primary human LECs. After confirming the low constitutive MHC I expression in human LECs, we observed a significant upregulation of MHC I across three chronic respiratory diseases: chronic obstructive pulmonary disease (COPD), idiopathic pulmonary fibrosis (IPF), and cystic fibrosis (CF). Additionally, we unveiled an unexpected sexual dimorphism in MHC I expression in both health and disease, with males exhibiting higher levels of MHC I under steady-state conditions. Gene expression analyses suggest that differential redox balance between sexes is instrumental in this dimorphism. Our study unveils the complex interplay between MHC I expression, sex, and respiratory diseases. Since, in other models, MHC I upregulation contributes to the development of immunopathologies, we propose that it might have a similar impact on chronic lung diseases.

**NEW & NOTEWORTHY:** This study shows that MHC I expression is very low in healthy LECs but escalates significantly in three chronic respiratory diseases, potentially contributing to disease progression. Furthermore, sex-specific divergences in LEC MHC I levels hint at distinct susceptibilities to chronic lung inflammation between males and females.

**Graphical abstract:** 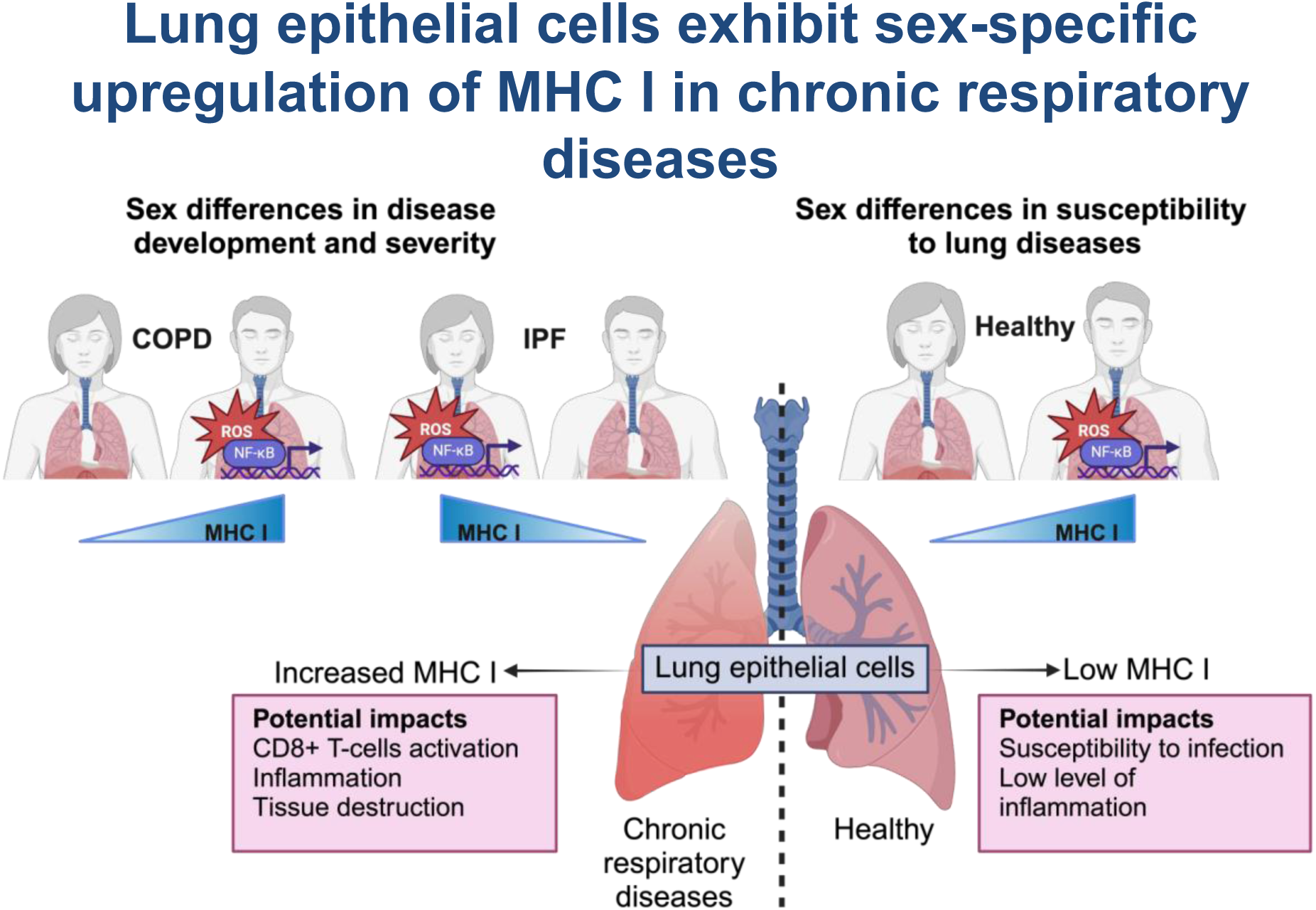

## INTRODUCTION

By scanning peptides presented by MHC I molecules, CD8+ T-cells perform immunosurveillance against infection and neoplastic transformation (1). MHC I expression also contributes to critical homeostatic functions within tissues, such as neuronal survival (2). The cellular machinery responsible for MHC I surface expression is complex, involving peptide generation by proteasomes, followed by translocation into the endoplasmic reticulum through Transporter associated with Antigen Processing 1 and 2 (TAP1 and 2) (3). Assisted by various chaperone proteins, peptides are loaded onto MHC I complexes, consisting of an MHC I heavy chain and beta-2-microglobulin (B2M). Once formed, the MHC-peptide complex is transported to the cell surface via the Golgi apparatus, ready for interaction with immune cells. MHC I expression is transcriptionally regulated at several levels (4). NOD-like receptor family caspase activation and recruitment domain containing 5 (NLRC5) regulates basal and induced expressions of MHC I components. The MHC I promoter also has binding sites for NF-κB and IFN sensitive response elements, allowing regulation by NF-κB signaling and IFN Regulatory Factors (IRF), respectively. MHC I is ubiquitously present in nearly all nucleated cells. However, recent studies have revealed heterogeneity in MHC I expression across tissues and cell types (5). Thus, quiescent tissue stem cells evade immune recognition by downregulating MHC I expression (6). On the other hand, MHC I overexpression is instrumental in various immunopathologies, including neurodegenerative diseases, myopathies, type I diabetes, and inflammatory bowel diseases (7–10). Hence, MHC I expression must be tightly regulated because of its pleiotropic effects.

Lung diseases, as reported annually by the World Health Organization, are among the leading causes of death worldwide. Chronic Obstructive Pulmonary Disease (COPD), Idiopathic Pulmonary Fibrosis (IPF), and Cystic Fibrosis (CF) are prominent examples of chronic diseases characterized by disrupted lung epithelium function, susceptibility to infections, and heightened inflammation (11–13). COPD and IPF are progressive and principally affect older adults with a smoking history. In contrast, CF is a genetic disease caused by mutations in the gene coding for the Cystic Fibrosis Transmembrane Conductance Regulator (CFTR) channel (14). These diseases are heterogeneous and exhibit sex dimorphisms in severity and pathogenesis (15–17). We reported that constitutive MHC expression is very low in mouse lung epithelial cells (LECs) but can be upregulated by inflammation driven by inhalation of LPS (18, 19).

We hypothesized that, as in mice, MHC I might be expressed at a low level in human LECs and be upregulated in chronic lung diseases. To address this hypothesis, we conducted comprehensive analyses using single-cell RNA sequencing (scRNA-seq) profiles obtained from lung parenchymal tissue of patients with COPD, IPF, CF, and non-diseased control donor lungs. We found a conspicuously low expression of MHC I in human LECs compared to other cell types in non-diseased lungs. LEC MHC I expression was significantly increased in COPD, IPF, and CF. Additionally, we unveiled a significant sex dimorphism in the expression of MHC I in healthy and diseased lungs. Modulation of MHC I expression could significantly impact the development of chronic lung diseases.

## MATERIALS AND METHODS

### Publicly available scRNA-seq Data Analyses

We analyzed 312,928 cells from 26 non-diseased, 18 COPD, and 28 IPF distal lung parenchyma explants publicly available (20). Sample processing and digestion procedures were described in the original paper. For CF donors, a Seurat object containing cells from 19 CF patients and 19 non-diseased control donors was downloaded (21). The sex, age, and smoking status of the donors were obtained from the original papers and are presented in Supplementary Table S1 (non-disease, COPD, and IPF donors) and Supplementary Table S2 (CF patients). Additionally, scRNA-seq data from intestinal epithelium were examined (22). All data were analyzed using Seurat V.4 (23). Quality control and cell annotation procedures followed the protocols outlined in the original studies (20–22).

### Differential Gene expression and GSEA analysis

Differential gene expression analysis was conducted using the Seurat function “FindMarkers”, comparing each parameter (sex or disease) one by one for each cell type of interest. We used a natural log fold-change threshold of 0.25, a minimum percentage of cells expressing a gene of 10%, and the Wilcoxon Rank Sum test. DEGs obtained for each comparison are shown in Supplementary Table S3. Gene Set Enrichment Analysis (GSEA) was performed using the “fgsea” function with Gene Ontology (GO) biological pathways terms as a gene set database from gsea-msigdb.org (24). We used the default parameters. Additionally, GO term enrichment analysis from DEGs (Figure 6) was conducted using https://www.pantherdb.org, using the GO biological process complete as a statistical overrepresentation test.

### Enrichment, purification, and cytocentrifugation of LECs from control donors and CF patients

Frozen LEC cytospin slides from six healthy and six CF patients (including three males and three females for each group) were provided by The Respiratory Tissue and Cell Biobank of CRCHUM, Montréal, Québec, according to the approved ethical protocol (Comité d’éthique de la Recherche Clinique de l’Université de Montréal, #2020-171: CERC-20-064-D and Comité de le recherche du CHUM/CRCHUM #08.063). All subjects signed informed consent; their demographics are described in Supplementary Table S4. Primary human airway (bronchial) and alveolar epithelial cells were obtained from explanted lungs from cystic fibrosis (CF) recipients during lung transplantation and biopsies from healthy (non-CF) donors. As previously detailed(13, 25–27), after dissection, bronchial tissues were rinsed and then incubated overnight at 4°C with MEM medium (Life Technologies, Burlington, QC, CA) supplemented with 7.5% NaHCO_3_ (Sigma-Aldrich, Saint-Louis, MO, USA), 2mM L-glutamine, 10 mM HEPES (Thermo-Fisher Scientific Inc., Waltham, MA, USA), 0.05 mg/ml gentamycin, 50 U/ml penicillin-streptomycin, 0.25 μg/ml Fungizone (Life Technologies) and containing 0.1% protease (from Streptomyces griseus; Sigma-Aldrich) and 10 μg/ml DNAse (Deoxyribonuclease I from bovine pancreas; Sigma-Aldrich). The protease-DNAse activity was then neutralized with FBS (Life Technologies), and bronchial epithelial cells were gently scraped off the remaining tissue. Red blood cells were removed from the cell suspension by treatment with ACK lysis buffer (0.1 mM NH_4_Cl, 10 μM KHCO_3_, 10 nM EDTA). Parenchymal lung tissues were rinsed with physiological saline solution, minced, digested with elastase (16 U/ml MEM, Worthington Biochemical, Lakewood, NJ, US, for 45 min), then finely chopped, and the resulting cell suspension was filtered. After centrifugation (300 g, 8 min), the cell pellet was resuspended in MEM. Alveolar epithelial cells were then purified using a differential adherence technique(28), which allows the discarding of remaining macrophages and fibroblasts attached to IgG-coated Petri dishes. Red blood cells were then removed by treatment with ACK lysis buffer (0.1 mM NH4Cl, 10 μM KHCO3, 10 nM EDTA). These steps allowed us to obtain a cell suspension enriched for alveolar epithelial cells (up to 86% of alveolar type II cells). Trypan blue staining of the post-IgG cell suspension confirmed >90% cell viability(29–33). After cell counting, the airway and alveolar cell suspensions were diluted to a density of 80,000 cells (in 200 μL PBS/slide) before cytocentrifugation (750 rpm, 5 min, Thermo Scientific Cytospin 4 Cytocentrifuge, Block Scientific, NY, US) onto glass slides).

### Immunofluorescence (IF) staining of human primary LECs

Cytospin slides from the primary airway and alveolar epithelial cells were fixed with 4% paraformaldehyde for eight minutes and subsequently blocked and treated with Wheat Germ Agglutinin (WGA Alexa Fluor 647, Invitrogen, Waltham, MA, catalog no. W32466) for 1 hour. Then, the samples were incubated overnight at 4°C with primary antibodies, including rabbit anti-pro-surfactant protein C (pro-SPC, Fisher Scientific, Waltham, MA, catalog no. AB3786), rabbit anti-cytokeratin 13 (CK13; Thermofisher, Waltham, MA, catalog no. MA5-32305), rabbit anti-βIV-tubulin (TUBB4; clone EPR1676, Abcam, Waltham, MA, catalog no. ab179509), rabbit anti-MUC5AC (Thermofisher, catalog no. MA5-12178), and mouse anti-HLA-ABC (clone W6/32; Biolegend, San Diego, CA, catalog no. 311402). Subsequently, sourced from Invitrogen, secondary antibodies anti-rabbit IgG Alexa Fluor 488 and anti-mouse Alexa Fluor 555 were applied. The pan anti-HLA-ABC antibody was omitted for a negative control of MHC I staining and replaced with an isotype. Imaging was performed using a Leica SP8 confocal microscope (Leica Microsystems, Wetzlar, Germany) at 400X magnification, and analysis was conducted using Image J2 software (version 2.3.0/1.53f, Rasband, W.S., National Institutes of Health, Bethesda, Maryland). Laser intensity settings were calibrated using negative controls. All donors were processed simultaneously and under the same conditions for each marker. No change in contrast or intensity was made.

### Statistical analysis

For scRNA-seq data, statistical significance was evaluated with a non-parametric Wilcoxon rank sum test using the Seurat function “find markers”. The type of comparisons and the *p* values are shown in the figure legends. Statistical significance was evaluated with an unpaired t-test using GraphPad Prism version 8 for immunofluorescence staining. *P* values are shown in the figure legends.

## RESULTS

### Human LECs express constitutively low amounts of MHC I transcripts

To investigate MHC I expression in non-diseased human LECs, we analyzed scRNA-seq data from distal lung parenchyma from 21 healthy non-smoking donors (20). We quantified the expression of the three classical human MHC I alleles (HLA-A, HLA-B, and HLA-C) and B2M across four major lung cell types: epithelial, lymphoid, myeloid, and fibroblastic. We selected cMonocytes as a reference because they are professional antigen-presenting cells (APCs). Relative to cMonocytes, MHC I expression was superior in other lympho-myeloid cells and inferior in fibroblasts and LECs (Figure 1A). LECs from the bronchioles (ciliated, club, basal, and goblet cells) and alveoli (alveolar type I (ATI), and alveolar type II (ATII)) expressed less MHC I than all other lung cell populations (Figure 1A). Next, we compared the expression of genes crucial for MHC I surface expression and their transcriptional regulation (Figure 1B-E). Key players in the peptide loading of MHC class I, such as *TAP1*, Calnexin (*CANX*), Calreticulin (*CALR*), and TAP binding protein (*TAPBP*) were expressed at low levels in ECs (Figure 1B). LECs also exhibit low amounts of constitutive (*PSMB5*, *PSMB6*, *PSMB7*) and immunoproteasome (*PSMB8*, *PSMB9*, *PSMB10*) subunits (Figure 1C) and diminished IFN-stimulated genes (ISGs) expression, including *NLRC5*, *STAT1*, *STAT2*, *IRF1*, *JAK1*, and *JAK2* (Figure 1D). While most of these genes are expressed at low levels in LECs, the differences reached statistical significance only relative to lymphoid cells (Figure 1B-D). However, LECs express significantly higher levels of genes encoding NF-κB inhibitors (*NFKBIA* and *NFKBIZ*) and lower levels of NF-κB subunits coding transcripts (*REL*, *RELB*, and *NFKB1*) (Figure 1E). Together, these results suggest that low constitutive MHC I may be explained by repression of NF-κB signaling.

**Figure 1.**
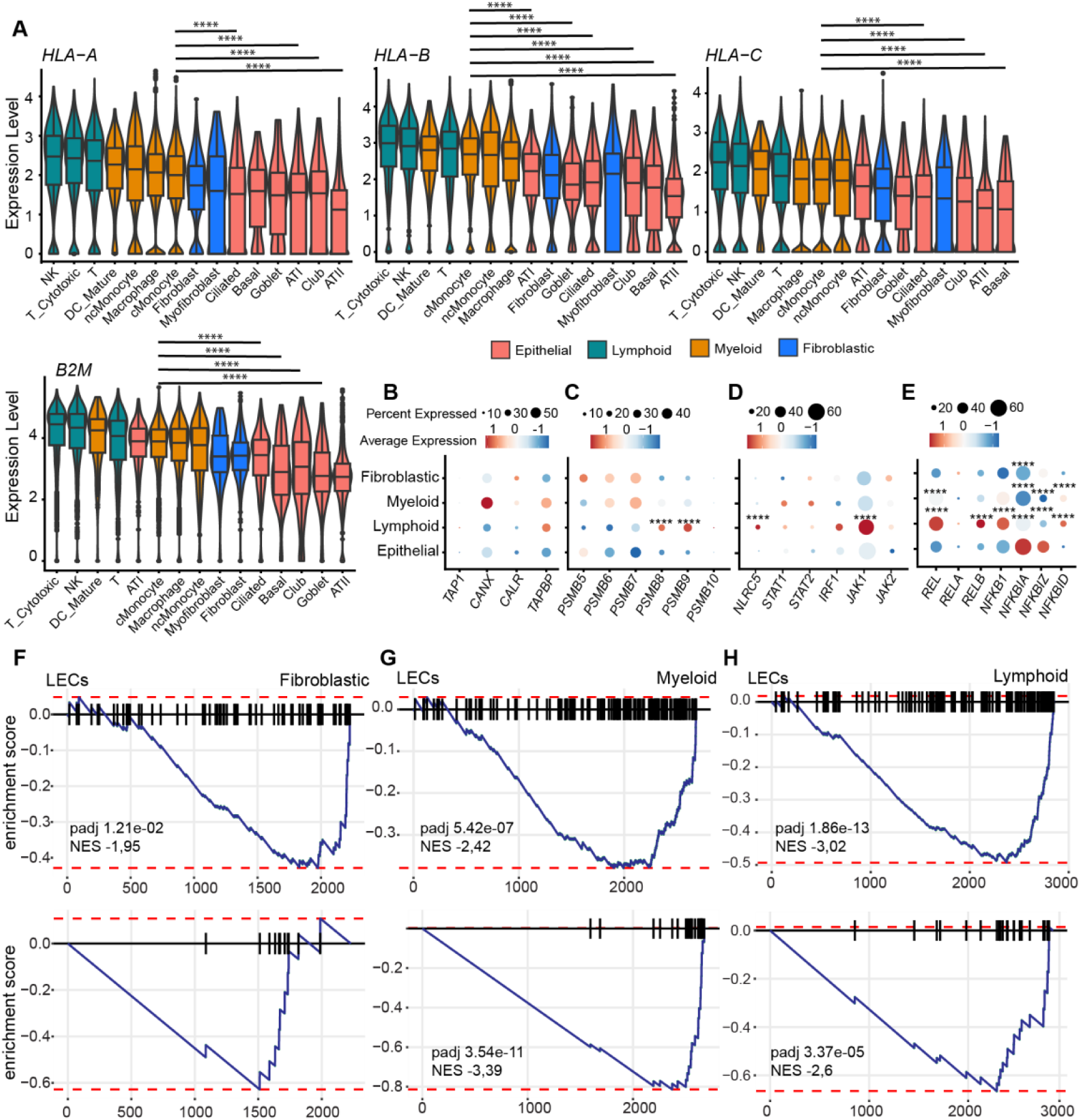
Human LECs constitutively express low amounts of MHC I transcripts. **A**) Comparison of MHC I alleles (*HLA-A*, *HLA-B*, *HLA-C*) and *B2M* transcript expression in scRNA-seq data from 21 healthy non-smoking donors, including 11 females and 10 males. The expression level corresponds to the feature counts for each cell divided by the total counts for that cell and multiplied by the scale factor (1000), then natural log-transformed using log1p. Statistical significance was assessed using a non-parametric Wilcoxon test to compare LECs and cMonocytes (**** *p* < 0.0001). Cell types are ordered from the highest MHC I expression to the lowest. **B-E**) Assessment of the average expression and the percentage of cells expressing PLC genes (B), constitutive and immunoproteasome subunits (C), ISGs involved in MHC I transcriptional regulation (D), and NF-κB signaling (transcription factors: *REL*, *RELA*, *RELB*, *NFKB1*; or inhibitors: *NFKBIA*, *NFKBIZ*, *NFKBID*) (E). Dot size reflects the percentage of cells with gene expression; color corresponds to the degree of expression. Statistical significance was assessed using a non-parametric Wilcoxon test (\**p* < 0.05, \*\**p* < 0.01, \*\*\**p* < 0.001, \*\*\*\**p* < 0.0001) comparing the average expression of each transcript in LECs related to each other cell types (lymphoid, myeloid, or fibroblastic cells). **F-H**) GSEA focusing on two GO biological pathways, “activation of immune response” (upper panels) and “antigen processing and presentation of peptide antigen” (lower panels) in LECs compared to fibroblastic cells (F), myeloid cells (G), and lymphoid cells (H). Normalized enrichment score (NES) and adjusted *p*-value (*p* adj) are shown on the graph when significant.

To pinpoint the key biological pathways that distinguish LECs from other lung cells, we conducted GSEA with DEGs obtained from the comparison between LECs and fibroblastic cells (Figure 1F), myeloid cells (Figure 1G), and lymphoid cells (Figure 1H). These analyses revealed a lower enrichment of the biological pathways: “activation of immune response” and “antigen processing and presentation of peptide antigen”, in LECs compared to the three other cell types (Figure 1F-H). Genes included in these pathways are described in Supplementary Table S5. These results reinforce that the low expression of genes linked to MHC I expression is a notable feature of LECs.

### Male LECs constitutively express more MHC I transcripts than females

Sex is one of the most significant sources of biological variability. Here, “sex” encompasses all biological differences between subjects genetically assigned as males and females. We conducted a sex comparative analysis of LECs obtained from non-diseased donors analyzed in Figure 1. Males exhibited significantly more MHC I transcripts than females within the Ciliated, Goblet, and ATII cell subsets (Figure 2A). This observation was validated at the protein level through IF staining of cytopsin slides of primary airway and alveolar epithelial cells (Figure 2B-C), revealing significantly higher MHC I fluorescence intensity in ATII (pro-SPC+ staining) and secretory cells (which include club and goblet cells, MUC5AC+ staining) from males compared to females. No significant difference was observed in ciliated cells at the protein level, probably because of the lower number of this cell population stained on each slide (data not shown). Sex differences in MHC I expression were not found in non-EC lung cells (Figure S1A) but were observed in certain types of ECs from the gut (Figure S1B). Hence, i) in the lung, only LECs show higher MHC I expression in males, and ii) this epithelial cell sex dimorphism is not limited to the lung.

**Figure 2.**
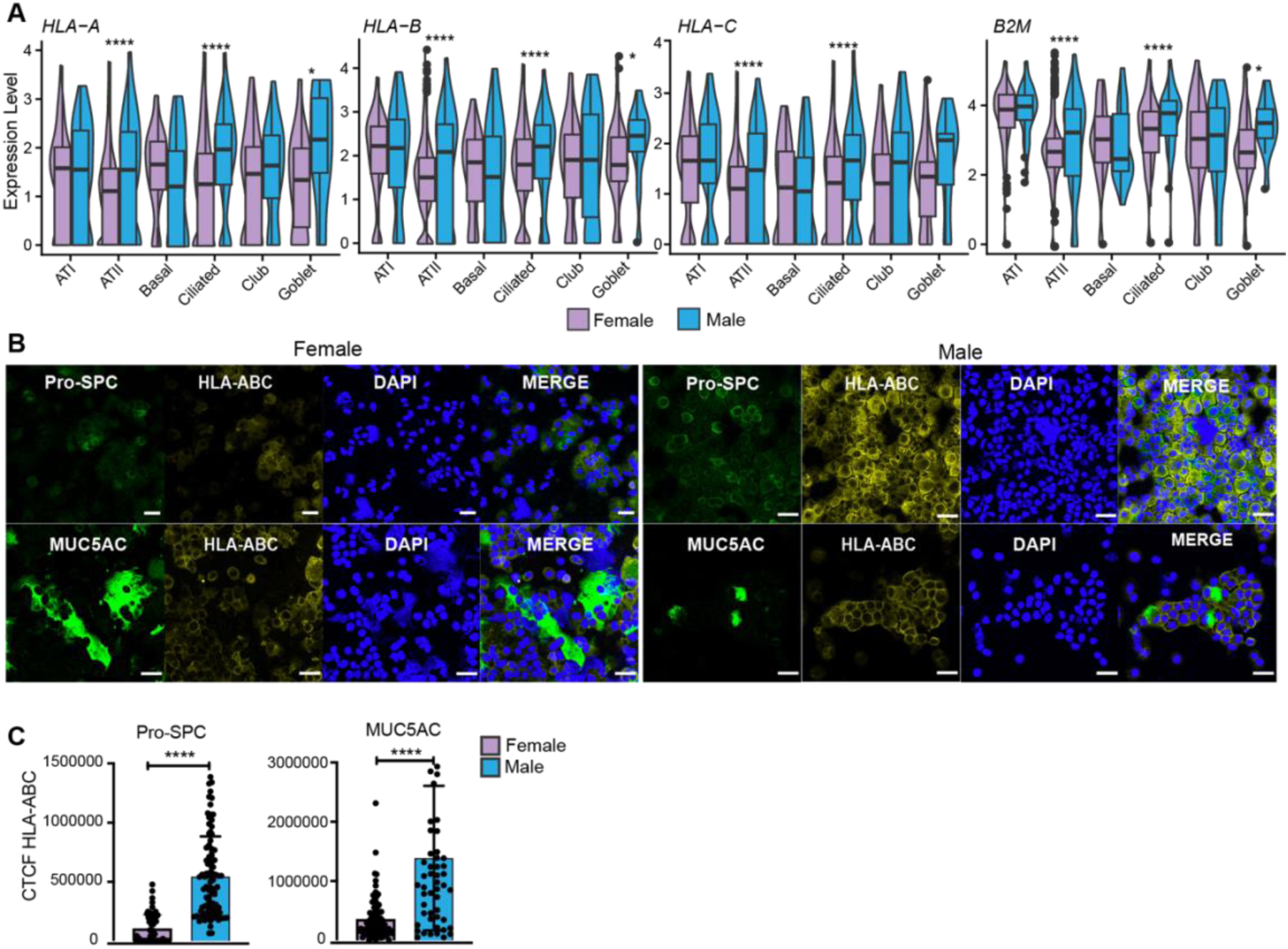
LECs express more MHC I in males than females in non-diseased lungs. **A**) Male (n=10) and female (n=11) comparison of MHC I alleles and B2M mRNAs expression levels in the healthy non-smoking donors analyzed in Figure 1. Statistical significance was assessed using a non-parametric Wilcoxon test comparing male and female LECs (\**p* < 0.05, \*\*\*\**p* < 0.0001). **B**) IF staining of human ATII (Pro-SPC+) and secretory (MUC5AC+) cells with pan anti-HLA-ABC antibody. Representative images for all staining conditions of non-diseased LECs from males (n=3) and females (n=3). Donors’ data are shown in Supplementary Table S4. Images were taken using a 40X objective. Scale bars, 30μm. All donors (males and females) were stained and acquired simultaneously in the same conditions for each marker. No changes in the intensity or contrast were made. **C**) Corrected Total Cell Fluorescence (CTCF) calculated using the formula: Integrated Density – (Area of selected cell x Mean fluorescence of background readings). The statistical significance of CTCF differences between males and females was determined using an unpaired t-test (\*\*\*\**p* < 0.0001). Graphs show the means of all cells acquired in each group ± SD.

We then compared the expression of genes involved in MHC I processing in ciliated and ATII cells (LECs showing the most significant sex dimorphism at the transcriptomic level) from males and females. Males expressed significantly more peptide-loading complex (PLC) genes (*CALR* and *CANX* in ciliated cells and *TAPBP* in ATII) than females (Figure 3A). Interestingly, the immunoproteasome subunit *PSMB9* expression was higher in males, while the constitutive subunit *PSMB5* expression was higher in females (Figure 3B). Although *NLRC5* and *STAT1* tended to be expressed at higher levels in males and JAK1 in females, these differences were insignificant (Figure 3C). Moreover, males expressed more *IFNGR*, with significant differences observed for *IFNGR2* in ciliated cells (Figure 3D). Of particular interest, molecules involved in the NF-κB pathway exhibited significant sex dimorphisms in their transcript levels. In ATII cells, NF-κB inhibitors were significantly more expressed in females than males (*NFKBIA* and *NFKBIZ*), whereas males generally expressed more NF-κB subunits in ATII and ciliated cells than females, which is significant in ciliated cells (*REL* and *NFKB1*) (Figure 3E). GSEA analysis revealed that males exhibited enrichment in biological pathways related to the immune response in ATII and ciliated cells. In contrast, female ciliated cells showed enrichment for pathways associated with their functional specificities (e.g., cilium movement and organization, which included *HYDIN*, *DNAH5*, *DNAH7*, and *DNAH9*) (Figure 3F). Regulation of reactive oxygen species (ROS) metabolism, which regulates NF-κB signaling and MHC I expression (34–37), ranked among the top five enriched pathways in male ATII cells (Figure 3F). The “Positive regulation of ROS metabolism” pathway was significantly enriched in male ATII and ciliated cells from males (Figure 3G). These results suggest that the sexual dimorphism in LEC MHC I expression is driven by differences in ROS production and, consequently, activation of the NF-κB pathway.

**Figure 3.**
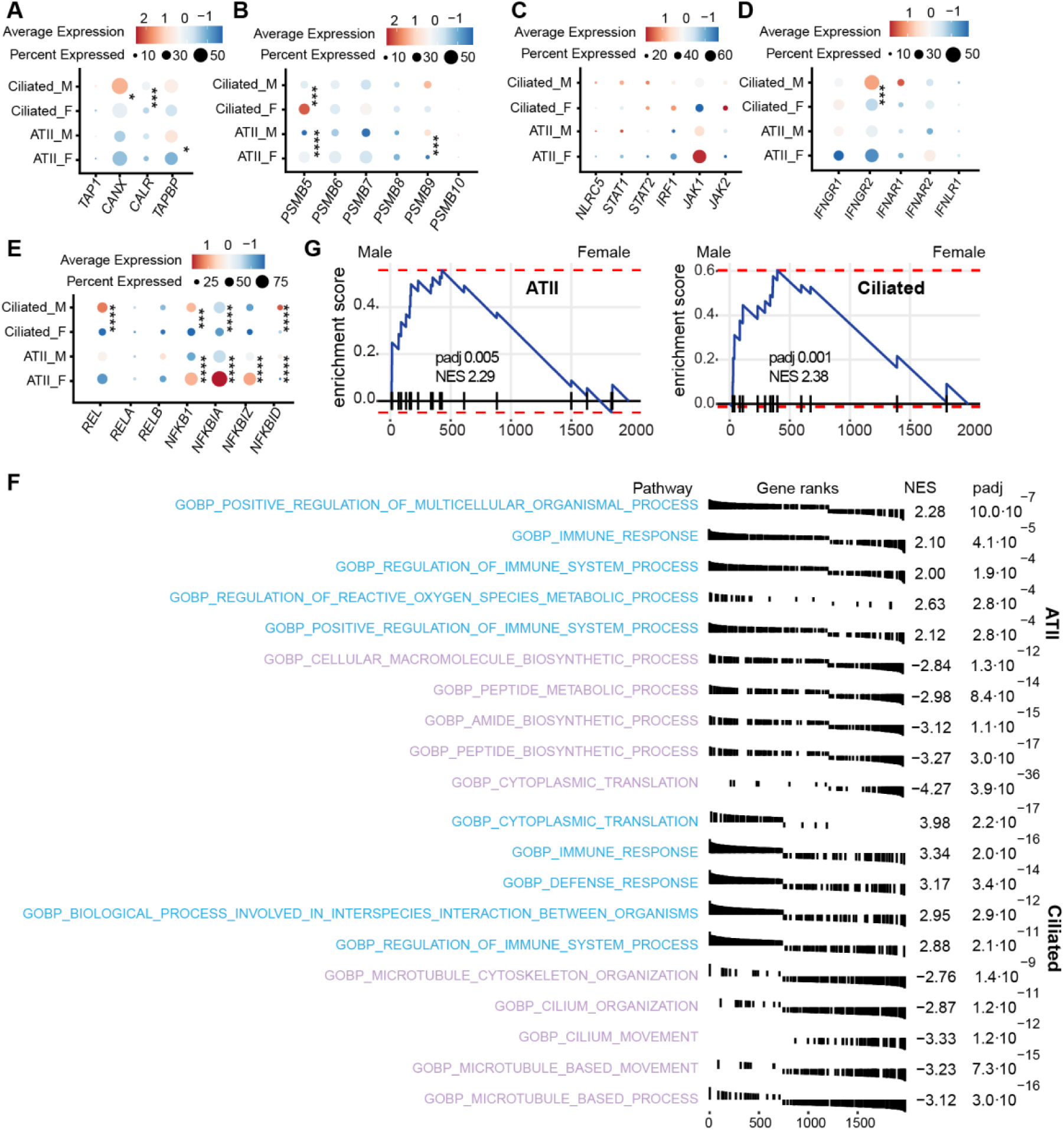
Male LECs express more transcripts contributing to MHC I expression than females. **A-E**) Percentage and average expression of ciliated and ATII cells expressing genes involved in the PLC (A), proteasome subunits (B), ISGs involved in MHC I transcriptional regulation (C), IFN receptors (D), and NF-κB signaling (E). Statistical significance was assessed using a non-parametric Wilcoxon test (\**p* < 0.05, \*\**p* < 0.01, \*\*\**p* < 0.001, \*\*\*\**p* < 0.0001) comparing the average expression of each transcript in male (n=10) versus female (n=11) healthy non-smoking donors for each cell type (ATII and ciliated cells). **F**) GSEA showing the top five enriched GO biological pathways in ATII and ciliated cells from males (blue) and females (purple). **G**) Enrichment score of the GO biological pathway “Positive regulation of reactive oxygen species metabolism” in male vs. female ATII and ciliated cells. NES and *p*adj are shown in the figure.

### Male LECs express higher amounts of genes involved in T-cell priming and cell death than females

In addition to MHC I presentation, priming of naïve CD8+ T-cells depends on the presence of costimulatory molecules on APCs (e.g., the more classical CD80 and CD86, as well as ICAM1, ICOS-L, and CD58) (38). Without costimulatory molecules, cells expressing MHC I can be recognized by primed effector CD8+ T-cells but cannot prime naïve CD8+ T-cells. The efficiency of CD8+ T-cell responses is further amplified by concurrent stimulation of CD4+ T-cells by peptides presented by MHC II molecules (e.g., HLA-DR). Among LECs, ATII are recognized as APCs and express APC-specific markers, including HLA-DR (39). A similar role has been proposed for ciliated cells (40). We examined the expression of genes coding for costimulatory molecules in ATII and ciliated cells from males and females. Males expressed significantly higher levels of *HLA-DR* and *CD86* than females (Figure 4A-B). On the other hand, females expressed more *ICAM1* (usually found in cells lacking *CD86*) (Figure 4B). We examined T-cell phenotypes to assess whether disparities in MHC I and costimulatory molecules may influence T-cell responses. Male T-cells showed higher late activation markers (*HLA-DR*) expression than female T-cells (Figure 4C). Furthermore, GSEA analysis performed in ATII and ciliated cells confirmed the enrichment of genes involved in “T-cell activation” in males compared to females (Figure 4D and Supplementary Table S5). Additionally, we observed an enrichment of genes related to the positive regulation of cell death in male ATII and ciliated cells compared to females (Figure 4E and Supplementary Table S5). Hence, alongside higher MHC I expression, ATII and ciliated cells from males overexpress genes involved in T-cell priming and cell death.

**Figure 4.**
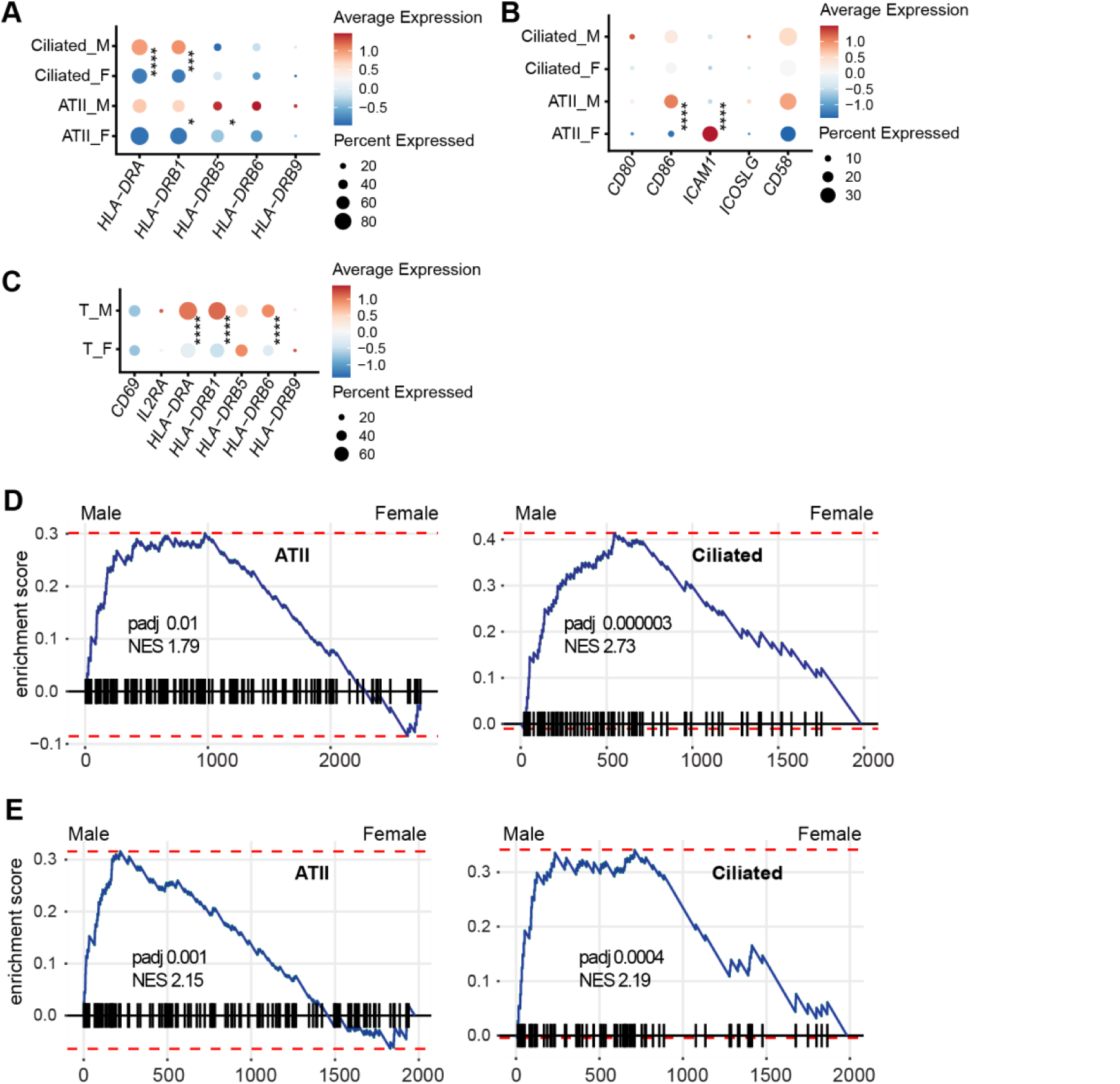
Male LECs express higher amounts of transcripts coding for T-cell activation and cell death markers than females. **A-B**) Percentage and average expression of ciliated and ATII cells expressing genes coding for HLA-DR (A) and costimulatory markers (B). **C**) Percentage and average expression of T-cell activation markers. Statistical significance was assessed using a non-parametric Wilcoxon test (\**p* < 0.05, \*\**p* < 0.01, \*\*\**p* < 0.001, \*\*\*\**p* < 0.0001) comparing the average expression of each transcript in male (n=10) versus female (n=11) healthy non-smoking donors for each cell type (ATII, ciliated and T-cell). **D-E**) Enrichment score of the GO biological pathways “T-cell activation” (D) and “positive regulation of cell death” (E) in ATII and ciliated cells from males and females. NES and *p*adj are shown in the figures.

### LECs increase MHC I expression in chronic respiratory diseases

In mice, inflammation triggered by LPS upregulates MHC I in LECs (19). We, therefore, wished to evaluate whether this would also be the case for LECs in patients with COPD and IPF. COPD is characterized by a robust inflammatory response involving CD8+ T-cells (11, 41). Conversely, IPF is marked by irreversible scarring of the distal lung, inflammation described as mild, and CD8+ T-cells correlate with the severity of the disease (12, 42). We initially compared the expression of the three MHC I alleles and *B2M* in LECs from COPD and IPF patients to that in non-diseased control donors. Each donor’s age and smoking status are detailed in Supplementary Table S1. We found a significant upregulation of MHC I expression in most LEC subsets from male and female COPD patients (Figure 5A-B). In IPF donors, the upregulation of MHC I transcripts was more restricted, particularly in males (panel A), where a significant increase in MHC I expression was observed only in Basal, Club, and ATI/II cells (Figure 5A-B). The increase in MHC I in COPD and IPF was less consistent among non-epithelial lung cell subsets. Among non-LECs, only lung fibroblasts showed a consistent rise in HLA-A, -B, and -C in males and females (Figure S2). Some leukocytes even exhibited a decrease in MHC I expression (Mature DCs in males with IPF; NK and T-cells in females with COPD or IPF). We conclude that the upregulation of MHC I in COPD and IPF is limited to non-hematolymphoid cells and is more conspicuous in LECs than fibroblasts.

**Figure 5.**
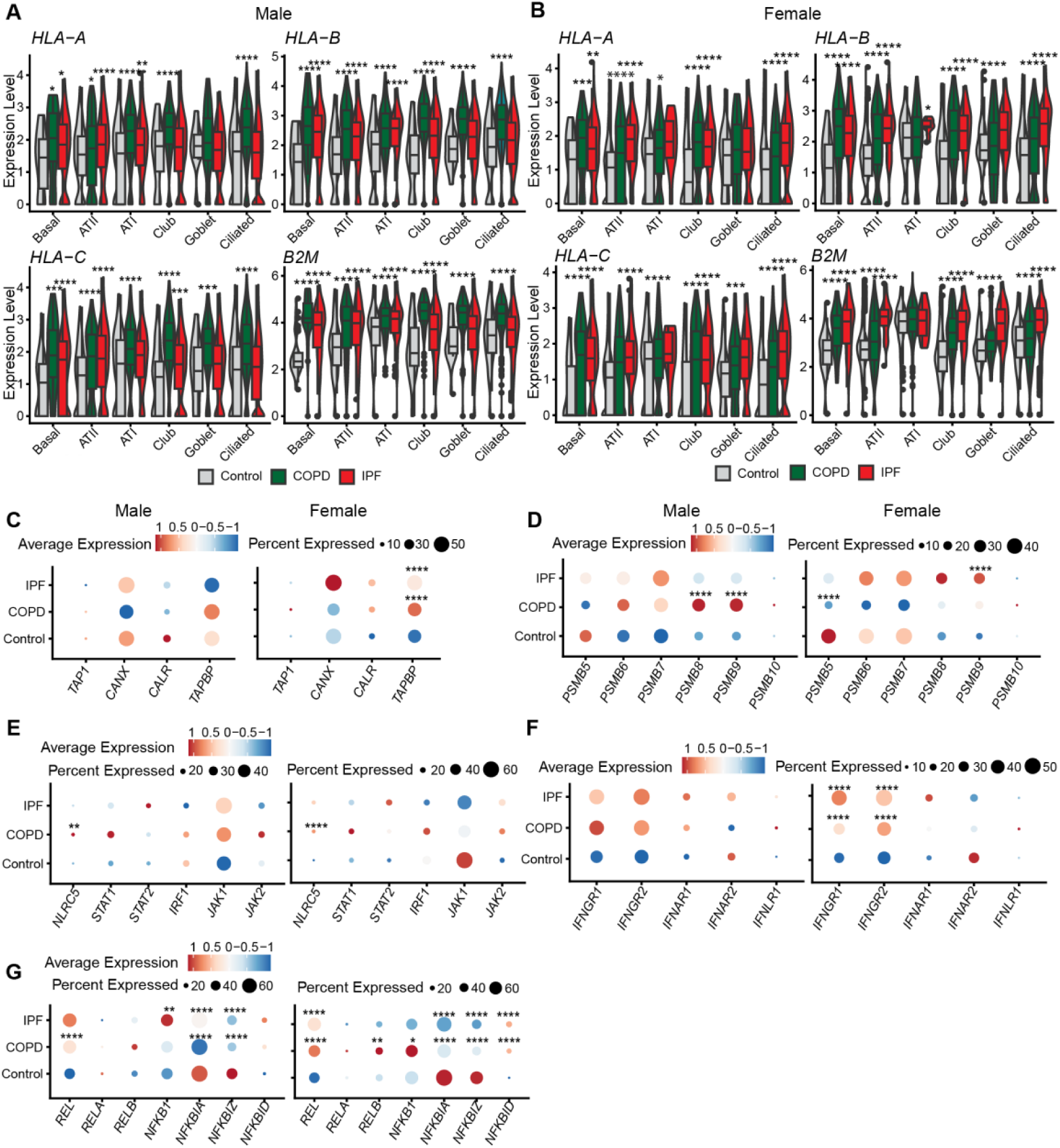
LECs upregulate MHC I-coding transcripts in chronic respiratory diseases. **A-B**) Expression levels of MHC I alleles and *B2M* transcripts in COPD (green), IPF (red), and non-diseased control donors (grey) in males (A) and females (B). The control group comprises 12 females and 14 males, the COPD group contains 9 females and 9 males, and the IPF group has 6 females and 22 males. All the groups included smokers and non-smokers. **C-G**) Average expression and percentage of LECs expressing genes involved in the PLC (C), proteasome subunits (D), ISGs involved in the transcriptomic regulation of MHC I (E), IFN receptors (F), and NF-κB signaling (G). All left panels display male data, and all right panels show female data. Statistical significance was assessed using a non-parametric Wilcoxon test (\**p* < 0.05, \*\**p* < 0.01, \*\*\**p* < 0.001, \*\*\*\**p* < 0.0001) comparing the average expression of each transcript in LECs from COPD or IPF patients versus non-diseased control donors. LECs analyzed in panels C to G contained merged LEC subtypes (ATI, ATII, basal, club, ciliated, and goblet cells), as in Figure 1.

We then looked at the expression of genes regulating the surface expression of MHC I (PLC and proteasome subunits) (Figure 5C-D) or its transcriptional regulation (IFN and NF-κB signaling) (Figure 5E-G). In COPD and IPF, male and female LECs showed no significant increase in PLC gene expression, except for *TAPBP* in females (Figure 5C). Expression of immunoproteasome subunits was increased in both sexes, with significant differences observed only in males with COPD (*PSMB8* and *9*) and females with IPF (*PSMB9*) (Figure 5D). NLRC5 expression was significantly higher in males and females with COPD, while other ISGs like *STAT1* and *IRF1* showed non-significant increases (Figure 5E). IPF patients exhibited no significant increase in ISG expression in either sex. Both males and females overexpressed IFNGR in COPD and IPF patients, but statistical significance was reached only in females for both diseases (Figure 5F). Additionally, both sexes exhibited increased expression of some NF-κB subunits (*REL*, *RELB*, or *NFKB1*) and decreased expression of NF-κB inhibitors (*NFKBIA*, *NFKBIZ*, or *NFKBID*), indicating NF-κB signaling activation in LECs in both pathologies (Figure 5G).

COPD and IPF are acquired diseases, prompting us to investigate whether a similar increase in MHC I expression occurs in genetic respiratory diseases like CF. We analyzed scRNA-seq data from CF distal lungs (bronchi and bronchioles), which were publicly available (21). As sex information was unavailable for all individuals (Supplementary Table S2), we merged male and female patients. Our analysis revealed increased HLA-B expression in ciliated, club, and goblet cells from patients with CF (Figure S3). We speculate that the rise in *HLA-A* and *C* expression might be attenuated due to the merging of males and females exhibiting sex dimorphisms in this analysis. Using IF staining of primary human alveolar and airway epithelial cells, we confirmed the increase in MHC I expression at the protein level in ATII and secretory cells from CF patients (Figure S4 and Supplementary Table S4).

Our findings demonstrate increased MHC I expression in LECs across three chronic respiratory diseases: COPD, IPF, and CF. This increase is likely due, at least in part, to IFN stimulation and NF-κB signaling in COPD and IPF.

### MHC I expression is differentially regulated in males and females with COPD and IPF

LECs showed higher constitutive MHC I levels in males than females (Figure 2). We therefore wished to investigate whether this difference persisted in the context of chronic respiratory diseases like COPD and IPF. We observed a significantly higher MHC I expression in males than females across all LEC subsets in the COPD donor group (Figure 6A). In IPF, the differences between sexes affected ATII and club cells and showed a different directionality (higher expression in females) (Figure 6B). Hence, males present a higher MHC I increase in COPD, whereas upregulation of MHC I in IPF is more marked in females (Figure S5A, B).

**Figure 6.**
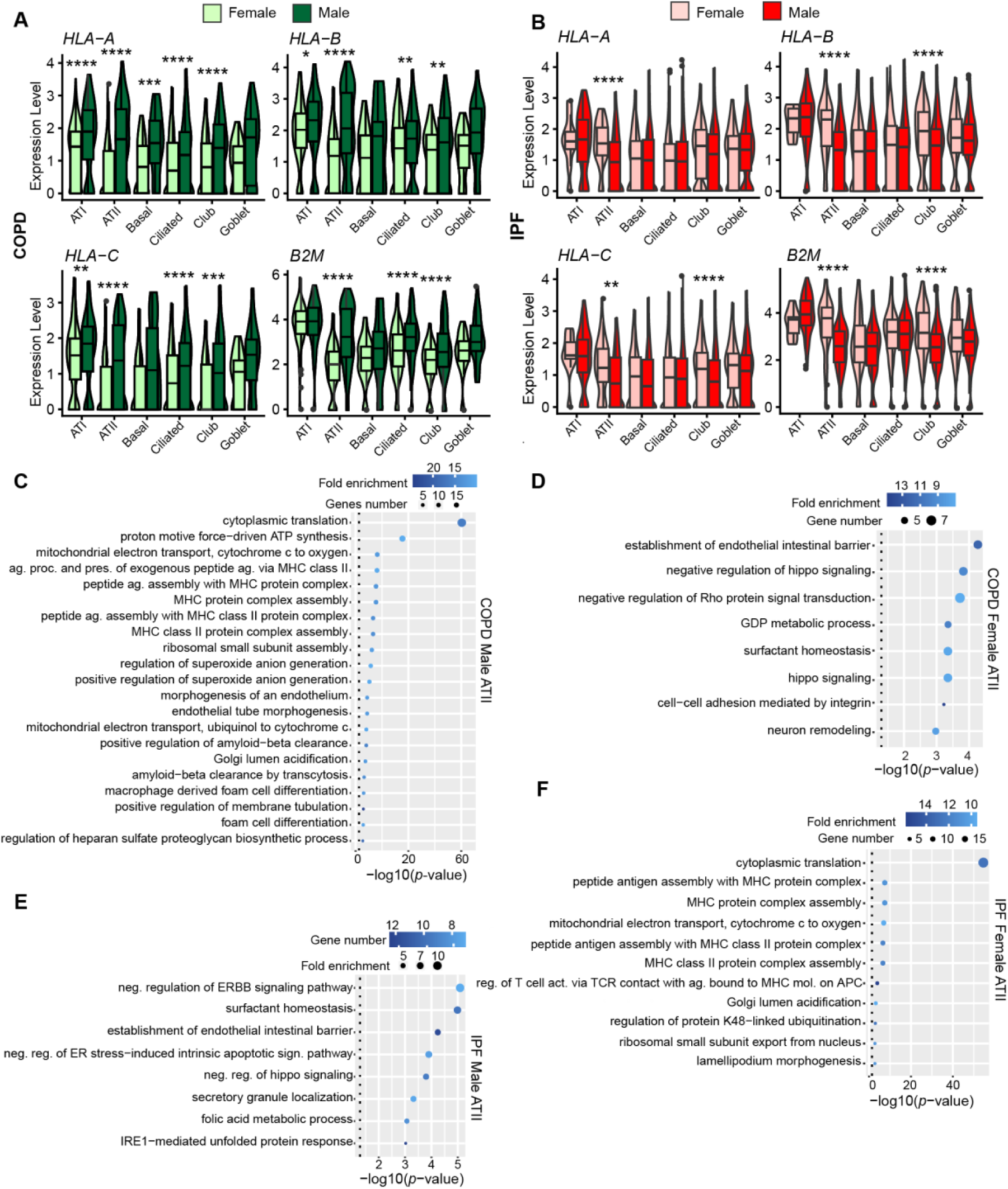
MHC I expression is differentially regulated in males vs. females with COPD and IPF. **A-B**) Expression levels of MHC I alleles and *B2M* in males (darker color) and females (lighter color) in COPD (n= 9 males and 9 females) (A) and IPF (n=22 males and 6 females) (B). Statistical significance was assessed using a non-parametric Wilcoxon test comparing the average expression of each transcript in males versus females (\**p* < 0.05, \*\**p* < 0.01, \*\*\**p* < 0.001, \*\*\*\**p* < 0.0001). **C-F**) GO biological pathway enriched in ATII cells from males (C) and females (D) with COPD and males (E) and females (F) with IPF. GO terms were selected according to their fold enrichment (superior to 10 for “COPD Male ATII” and “IPF Female ATII” and superior to 7 for “COPD Female ATII” and “IPF Male ATII”).

To understand the differential regulation of MHC I between males and females in COPD and IPF, we analyzed DEGs from ATII cells. We focused on ATII cells since the sex differences in this cell type were most consistent and striking throughout our study. Biological pathway analysis revealed that in COPD patients, male ATII cells exhibited enrichment in pathways related to antigen presentation and ROS generation (e.g., superoxide anion generation, mitochondrial respiration, (Figure 6C and S5C-F). Conversely, female ATII cells showed enrichment in their specific function-related pathways (e.g., surfactant homeostasis, including *SFTPD* and *ABCA3* genes, and regulation of hippo signaling, including *YAP1* and *SAV1* genes) (Figure 6D). In IPF, male ATII cells showed enrichment of genes involved in surfactant homeostasis and regulation of hippo signaling pathways (Figure 6E), and female ATII in pathways related to MHC assembly and mitochondrial respiration (Figure 6F and S5C-F). DEGs associated with ROS generation and mitochondrial respiration correlated with HLA-B expression in males and females, emphasizing their putative role in regulating MHC I expression (Figure S5C-F). These findings show that inherent differences in ROS levels between males and females correlate with MHC I expression in LECs at steady state and in chronic respiratory diseases.

To assess the potential impact of these disparities on T-cell activation in COPD and IPF, we examined genes involved in T-cell priming in ATII and ciliated cells. In COPD, male ATII cells exhibited elevated levels of HLA-DR, CD86, and CD58 (Figure 7A-B). Conversely, in IPF, females displayed increased HLA-DR and CD86 levels, with males exhibiting higher ICAM1 expression (Figure 7A-B). Furthermore, male T-cells exhibited significantly higher levels of the late activation marker HLA-DR. Besides, females with COPD showed elevated levels of the early activation marker CD69 (Figure 7C). IPF donors showed no significant differences in T-cell activation markers (Figure 7C). Finally, GSEA performed in ATII from males and females showed T-cell activation (Figure 7D) and cell death-related gene (Figure 7E) enrichment in males with COPD and females with IPF (Supplementary Table S5).

**Figure 7.**
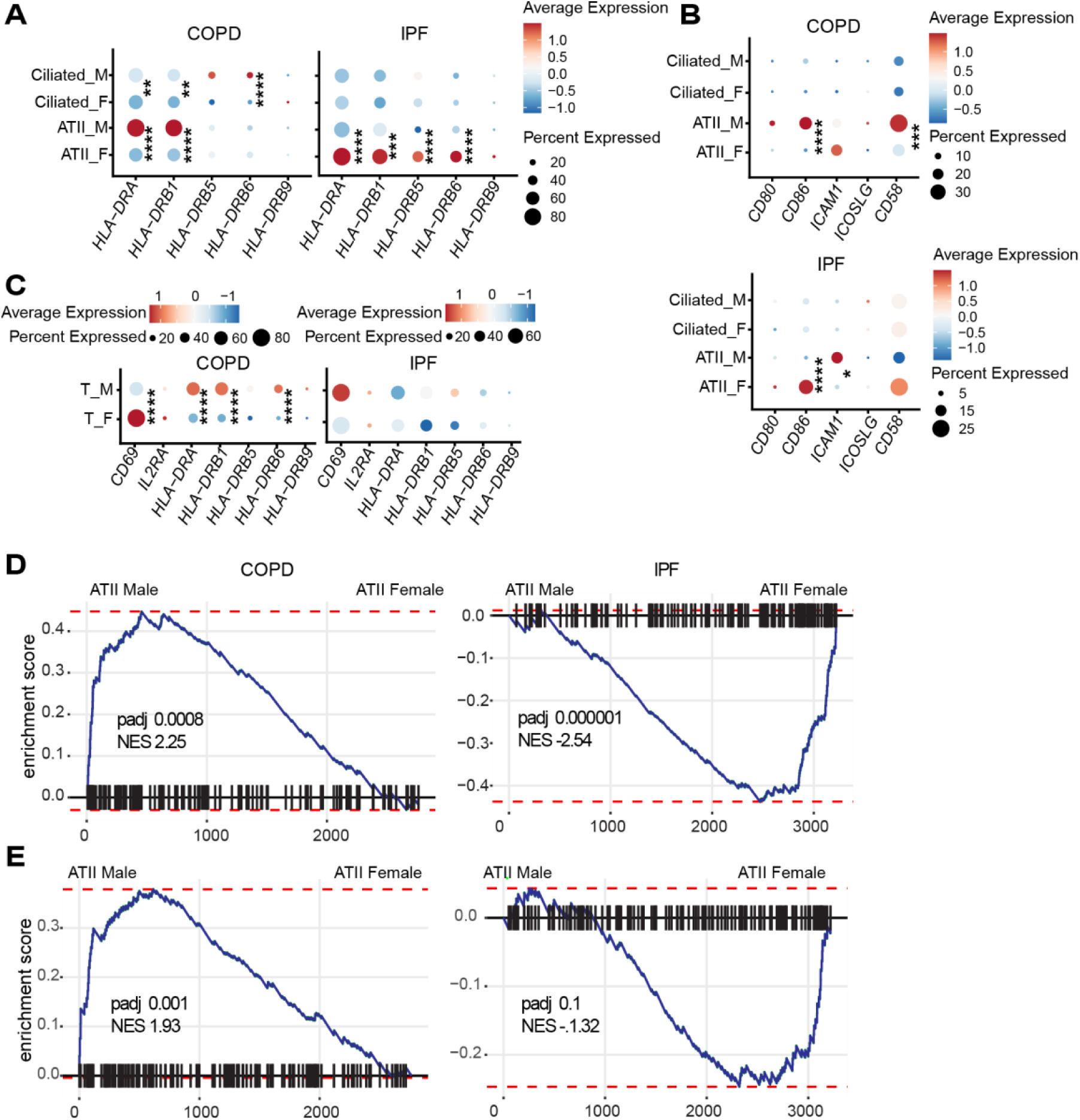
LECs differentially regulate transcripts coding for T-cell activation and cell death markers in males vs. females with COPD and IPF. **A-B)** Average expression and percentage of ATII and ciliated cells expressing *HLA-DR* (A) and costimulatory molecules (B) in males and females with COPD (n= 9 males and 9 females) and IPF (n=22 males and 6 females). **C**) Average expression and percentage of T-cell expressing T-cell activation markers with COPD and IPF. Statistical significance was assessed using a non-parametric Wilcoxon test (\**p* < 0.05, \*\**p* < 0.01, \*\*\**p* < 0.001, \*\*\*\**p* < 0.0001) comparing the average expression of each transcript between males and females in each cell type (ATII, ciliated, T-cell). **D-E**) Enrichment score of the GO biological pathways “T-cell activation” (D) and “positive regulation of cell death” (E) in ATII cells from male compared to female patients with COPD or IPF. NES and *p*adj are shown in the figures.

These findings underscore significant sex-based differences in the expression of genes related to MHC I, costimulatory molecules, and cell death markers among non-diseased individuals and those with COPD and IPF, with variations specific to each disease. These variations also extend to genes coding for T-cell activation markers.

## DISCUSSION

The lung faces constant exposure to airborne pathogens and particles, rendering it susceptible to inflammation-induced damage. Therefore, the lung must maintain a delicate balance: rapid clearance of pathogens without undue harm to LECs. Accordingly, dysregulated tissue-immune interactions are the hallmark of diverse chronic lung diseases (43–45). This study delves into the factors governing MHC I expression and the linked susceptibility to CD8+ T-cell mediated damage in healthy and diseased LECs.

MHC I expression is transcriptionally regulated by ISGs (particularly NLRC5) and NF-κB (4). The basal expression of MHC I is affected primarily by ISGs in some cell types (e.g., neurons) and by NF-κB in other cell types (e.g., astrocytes and oligodendrocytes) (46). We report that, like mouse LECs (19), human LECs constitutively express very low levels of MHC I. Also, human LECs express low amounts of NLRC5, other ISGs, and various genes responsible for MHC processing. However, compared to other lung cell types, MHC I expression in LECs correlated more strongly with decreased NF-κB-related gene expression.

Notably, we found that in chronic lung diseases, MHC I upregulation was strong in LECs, mild in fibroblasts, and absent in lympho-myeloid cells. Thus, chronic lung inflammation modulated MHC I expression primarily in the cell type with the lowest basal expression (LECs). This is reminiscent of other studies where disease correlated with MHC I upregulation in cell types with low basal expression, such as neurons, myocytes, and enterocytes (7, 8, 10). This suggests that fluctuations in MHC I levels in chronic inflammatory settings predominantly affect cells with low basal expression. Global MHC upregulation increases the abundance and diversity of MHC-associated peptides and could pave the way to autoimmunity (47, 48). Coherent with this idea, the lungs of patients with COPD and IPF commonly exhibit increased CD8+ T-cells (11, 12, 41, 49). Mechanistically, our transcriptomic analyses of LECs revealed an activation of NF-κB signaling in COPD and IPF. Upregulation of *NLRC5* and *IFNGR* was limited to COPD, where inflammation is more intense than in IPF.

LPS-induced MHC I upregulation in mice correlates with the downregulation of genes crucial for LEC functions, such as cell differentiation (ATII) and cilium movement (ciliated cells) (19). Likewise, in human LECs, we observed an enrichment of genes implicated in epithelial cell functions in conditions where MHC I expression was low. ATII from females in COPD and males in IPF are enriched in genes involved in hippo signaling (*YAP1*, crucial for their differentiation in ATI) (50) and surfactant homeostasis (*SFTPD* and *ABCA3*). In contrast, ciliated from healthy females are enriched in genes involved in cilium movement (*HYDIN*, *DNAH5*, *DNAH7*, and *DNAH9*). Thus, overexpression of MHC I may be a marker of defective epithelial cell functionality. The mechanistic underpinnings of this correlation have yet to be elucidated.

An unexpected finding from our study is the higher expression of MHC I observed in male ATII, goblet, and ciliated cells compared to their female counterparts. The superior MHC I expression in male LECs is accompanied by the upregulation of genes involved in T-cell priming. In non-diseased LECs, this sex dimorphism aligns with elevated levels of NF-κB subunits in males and NF-κB inhibitors in females. Various stimuli, such as cytokines, hormones, and cellular stress, can modulate NF-κB signaling (51, 52). Cellular stress, specifically ROS production, emerges as one of the top five enriched biological pathways in male ATII and ciliated cells. The role of ROS in regulating MHC I expression has been extensively studied in DCs and tumor cells (35–37). Considering the enrichment of genes related to ROS production in male LECs and the known role of ROS in regulating MHC I expression, we posit that sex differences in the redox balance of the lung contribute to the observed sex-specific regulation of MHC I expression in healthy LECs. ROS have been found to elevate the expression of PLC and immunoproteasome subunits (53). In accordance with this, we found that these genes are expressed at higher levels in male LECs than in females. Sex hormones play pleiotropic roles in physiological differences between males and females. Notably, estrogens have been shown to inhibit NF-κB signaling (54), the master regulator of MHC I expression in human LECs. Additionally, estrogens suppress ROS production by modulating antioxidant enzyme activity, whereas testosterone exhibits pro-oxidant properties (55–58). As estrogen receptors’ expression in the lung epithelium is well-documented (59), we propose that sex hormones contribute to the sex-specific regulation of MHC I in LECs by modulating ROS production and NF-κB signaling pathways.

To our knowledge, sex dimorphism in the expression of MHC I-related genes has been reported in two contexts: the aging brain, where aging females express higher levels of MHC I than males, and in peripheral blood mononuclear cells, where immunoproteasome activity is lower in females compared to males (60, 61). These two studies raised the possibility that dimorphisms of MHC I expression play a role in the differences between males and females in pathological contexts. Also, they indicate that inter-sex differences in MHC I regulation may not have the same directionality in all cell types. Chronic respiratory diseases exhibit notable sex differences in severity and pathophysiology (15, 17, 62). We documented that, in COPD, males show a more pronounced upregulation of MHC I and antigen presentation-related genes than females. The opposite trend is observed in IPF. Interestingly, in both diseases, the sex exhibiting higher MHC I expression shows enrichment in genes associated with mitochondrial respiration and ROS production. The involvement of redox balance in sex dimorphisms related to diseases has been demonstrated across various contexts, including COPD (16, 57). Furthermore, oxidative stress plays a significant role in COPD, CF, and IPF (63–66). The most parsimonious model to explain these data is that sex hormones influence the sex-specific regulation of MHC I expression by the redox balance and contribute to the sex differences in chronic respiratory diseases.

We acknowledge that our study presents several limitations. First, it relies mainly on transcriptomic data. The limited availability of human lung tissue samples has posed challenges for conducting further protein analysis in patients with COPD and IPF. Significant age differences exist between the control and COPD/IPF groups, reflecting the typical elderly population affected by these conditions (females median age: 48 for control, 60 for COPD, and 66.5 for IPF; males median age: 43.25 for control, 65 for COPD, and 66.5 for IPF). We chose not to exclude younger donors from the control group to avoid compromising the sample size. Finally, the functional implications of MHC I overexpression on lung epithelium have yet to be worked out. Future investigations, encompassing murine models and human subjects, should strive to elucidate the active role of MHC I in LEC biology, which we believe could be crucial to their function.

In summary, our study offers novel insights into the regulation of MHC I in both healthy and diseased human lungs. It illustrates how MHC I expression and the immune context, in general, are modulated by multiple factors that vary from one cell type to another in a sex-specific manner. Furthermore, our work raises several questions warranting further investigation. Two may be a priority. First, does MHC I upregulation in LECs lead to immune-mediated damage in chronic inflammatory lung diseases? Second, does the low basal MHC I expression in LECs hamper cancer immunosurveillance?

## DATA AVAILABILITY

Raw scRNA-seq data from control, COPD, and IPF donor lungs are available on GEO under the accession number **GSE136831**: https://www.ncbi.nlm.nih.gov/geo/query/acc.cgi?acc=GSE136831

Raw scRNA-seq data from control and CF donor lungs are available on GEO under the accession number **GSE150674**: https://www.ncbi.nlm.nih.gov/geo/query/acc.cgi?acc=GSE150674

Raw scRNA-seq data from non-diseased intestines are available on the Broad Institute Single Cell Portal under accession number **SCP259**: https://portals.broadinstitute.org/single_cell/study/SCP259

## SUPPLEMENTAL MATERIAL

**Supplementary tables S1-S5 and figures S1-S5 are available here:** https://figshare.com/s/69076bdead66c8e5f12d

## ACKNOWLEDGMENTS

We thank the dedicated staff at the IRIC facilities, including the genomics, bioinformatics, microscopy, and histology research units, for their valuable assistance and insightful discussions. Special thanks are owed to the Quebec Respiratory Tissue and Cell Biobank of CR-CHUM for generously providing the human primary cells crucial for the immunofluorescence staining analyses reported in this paper. The graphical abstract was created with BioRender.com.

## AUTHOR CONTRIBUTIONS

JM, SB, and CP conceptualized the study and drafted the initial manuscript. JM conducted the experiments and performed the data analysis. JM and SB discussed the findings and prepared the figures. EB and DA assisted in designing, analyzing, and discussing the immunofluorescence analysis and provided the human cytospin slides from healthy subjects and cystic fibrosis patients. CP, SB, and EB supervised the study. All authors reviewed the manuscript.

